# Data-driven optimization yielded a highly-efficient CRISPR/Cas9 system for gene editing in Arabidopsis

**DOI:** 10.1101/2023.10.09.561629

**Authors:** Haiying Geng, Yifan Wang, Yupu Xu, Yu Zhang, Ershang Han, Yuming Peng, Zhenxing Geng, Yi Liu, Yue Qin, Shisong Ma

## Abstract

The CRISPR/Cas9 technology is widely used for plant gene editing. In Arabidopsis, the early CRISPR/Cas9 systems using constitutive promoters to drive *Cas9* expression usually have low heritable mutation efficiencies in the T1 generation. Germ cell-specific or cell division-specific promoters, like *EC1*.*2en-EC1*.*1p fusion, YAO* and *CDC45*, have been used as alternatives to direct *Cas9* expression, but the efficiency of getting nonchimeric mutations is still not high. To further improve gene editing efficiency, we used gene co-expression network analysis to identify *NUC1* as a candidate promoter for driving *Cas9* expression. *NUC1* has expression patterns similar to *YAO*, but expresses at a much higher level. We constructed a CRISPR/Cas9 system pHY07 that uses the *NUC1* promoter to drive *Cas9* and carries the mCherry protein as a fluorescent marker for selecting transgenic seeds. Using this system to edit the *GL2* gene, we obtained apparently nonchimeric mutations in 55% of the T1 transgenic plants. Among the Cas9-free T2 plants 99% harbored mutations that are stably inherited from the previous generation. Therefore, our system exhibited extremely high editing efficiency, and through fluorescent screening of transgenic seeds, it become easy to obtain Cas9-free and stable genetic mutants in the T2 generation.

## Introduction

In recent years, targeted gene editing based on CRISPR (clustered regularly interspaced short palindromic repeats)/Cas (CRISPR associated) in the bacterial immune system has become a powerful gene knockout technology. The CRISPR locus is transcribed into non-coding guide RNA, which forms a complex with Cas protein to mediate the cleavage of complementary invading DNA [1, 2]. Getting gene knockout materials through CRISPR/Cas9 gene editing methods, especially when there are no suitable mutants, is important to study the biological functions of genes in plants. At present, the CRISPR/Cas9 system has been applied to a variety of plants, such as Arabidopsis, rice, and tobacco [3-9].

In Arabidopsis, CRISPR/Cas9 gene editing is usually achieved by generating transgenic plants via *Agrobacterium*-mediated transformation using the floral dip method, which inserts CRISPR/Cas9 cassettes, containing the *Cas9* gene and a chimeric single guide RNA (sgRNA) sequence, into the genome of Arabidopsis egg cells or embryonic stem cells [10, 11]. In the early CRISPR/Cas9 systems, the constitutive Cauliflower Mosaic Virus (CaMV) 35S promoter was used to drive the expression of *Cas9*. However, the editing efficiency is very low, and most of the editing events in the T1 generation generate somatic mutations that do not inherit to the next generation. This is mainly because that, although CaMV 35S promoter is strongly expressed in vegetative tissues, its expression level is low in plant germ cells, but only mutations that occur in germ cells can be passed on to the next generation [6, 9, 10, 12]. Later researches showed that the editing efficiency of CRISPR/Cas9 systems are greatly affected by *Cas9* expression timing and tissue specificity. Several studies have used alternative promoters to drive *Cas9* expression to improve gene editing efficiency, such as egg cell specific *EC1*.*2en-EC1*.*1p* fusion promoter, cell division-specific *YAO* promoter, meiosis-specific *CDC45* promoter, dividing tissue-specific *ICU2* promoter, and constitutive *RPS5a* promoter that is expressed from an early embryonic stage [13-17]. Using these modified systems, heritable nonchimeric mutant plants can be obtained in the T1 generation, but the efficiency is still not high. The results of Feng et al. showed that, when using pCDC45-Cas9 and pYAO-Cas9 to edit the *GL2* gene, the efficiencies of obtaining nonchimeric mutants in the T1 generation were 4.4% and 10%, respectively [18]. Generally, two or three generations of screening are required to obtain single-gene homozygous mutants. The entire process is time-consuming and laborious. The efficiency of editing multiple genes is even lower. Therefore, a more efficient CRISPR/Cas9 editing system is needed to speed up the acquisition of gene knockout materials.

Here, we employed a data-driven strategy to identify potential gene promoters for improving CRISPR/Cas9 editing efficiency. Via gene co-expression network analysis, we discovered the *NUC1* gene promoter might be a good candidate. The *NUC1* gene has similar expression patterns to the *YAO* gene, but its expression level is much higher than *YAO*. We used the *NUC1* promoter to drive the expression of *Cas9* and constructed a CRISPR/Cas9 gene editing system named pHY07. Using this system to target the Arabidopsis *GL2* gene, we obtained apparently nonchimeric mutations in 55% of the T1 transgenic plants. Our system also contains the *mCherry* fluorescent marker gene driven by a seed specific promoter *At2S3*, which enabled us to visually select Cas9-free seeds from the T1 plants [19, 20]. Among the Cas9-free T2 plants, 99% harbored mutations that were stably inherited from the T1 generation. Similar results were also obtained when editing the *TRY* and *CPC* genes or multiplex editing the *GATA1* and *GATA5* genes simultaneously. Our system exhibited extremely high editing efficiency, and through fluorescent screening it become easy to obtain Cas9-free stable mutants in the T2 generation.

## Results

### Discovery of an efficient promoter and construction of a modified CRISPR/Cas9 system

In this study, we sought to improve the gene editing efficiency of CRISPR/Cas9 by selecting better promoters for driving *Cas9* expression. We reasoned that the ideal promoters should have similar expression patterns to those used previously for generating heritable mutations, so their expression timing and tissue specificity are right for the task. However, they should express at much higher levels than those used before, such that their editing efficiency will become higher. Previously, we have used large-scaled microarray data to construct a gene co-expression network based on the graphical Gaussian model called AtGGM2014, from which gene co-expression modules were identified [21]. The *YAO* gene, reported by Yan *et al*. before [15], was found to be within Module #21 (Fig. 1a). Theoretically, all the genes within this module should have similar expression patterns to *YAO* and can be used as alternative promoters to drive *Cas9* expression. We used large-scaled publically available RNA-Seq datasets to compare the expression levels of all the genes within the module, and *NUC1* (*PARL1*) was the most highly expressed one among them (Fig. 1a). Previous studies have shown that *NUC1* is mainly expressed in regions related to embryonic development and cell division, which is similar to *YAO* [15, 22]. Our analysis showed that *NUC1* is expressed 16 times higher than *YAO*. The *NUC1* promoter was then used to drive the expression of *zCas9* (*Zea mays* codon-optimized *Cas9*) to construct a modified CRISPR/Cas9 system named pHY07 (Fig. 1b). The *zCas9* sequence, together with a *Pisum sativum* rbcS E9 terminator, was obtained from a plant CRISPR/Cas9 vector pHEE401 published previously [14, 23]. The modified system also contains the *mCherry* gene driven by a seed specific promoter *At2S3*, which serves as a fluorescent marker for selecting transgenic seeds [19, 20].

**Figure 1.**
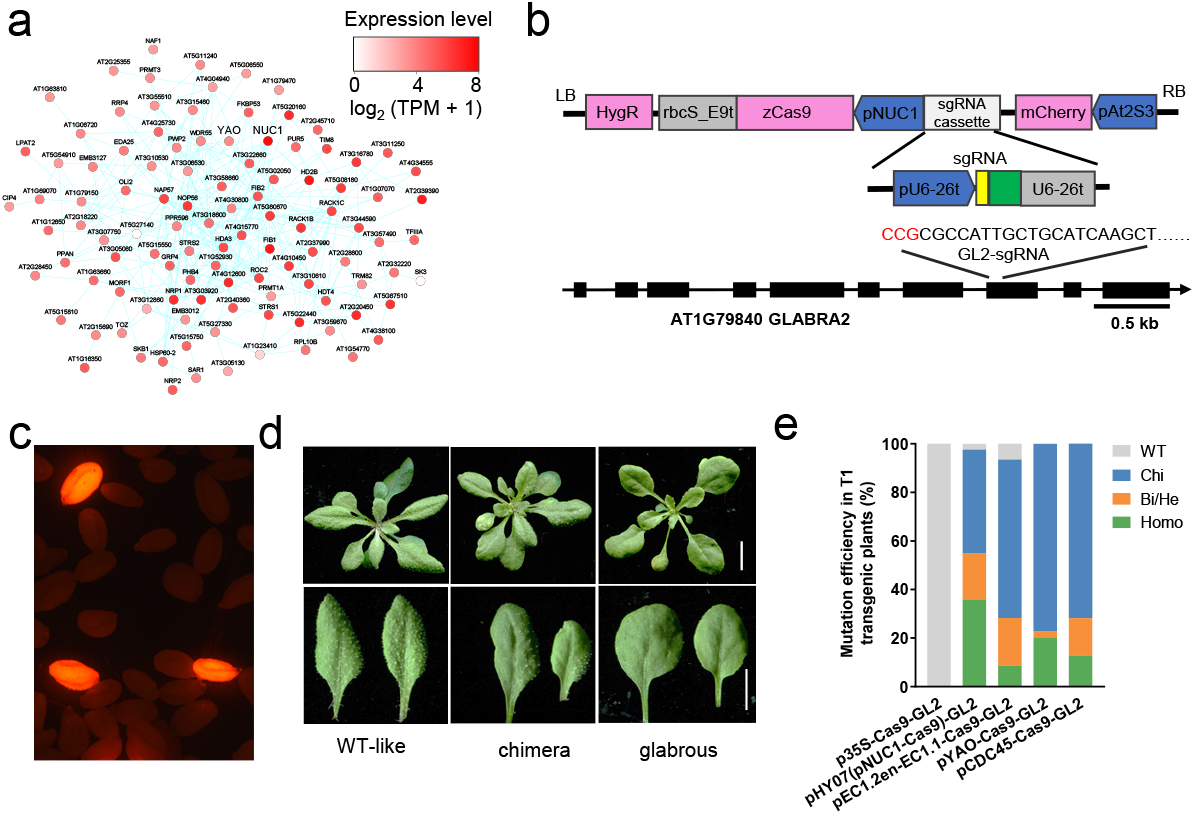
Gene editing efficiency of an optimized CRISPR/Cas9 system pHY07 in T1 generation transgenic Arabidopsis. **a**, Identification of *NUC1* as a candidate gene promoter for improving CRISPR/Cas9 gene editing efficiency. A gene co-expression Module #21 from an Arabidopsis gene co-expression network AtGGM2014 is shown. Nodes represent genes, colors indicate gene expression level, and co-expressed gene pairs are connected by edges. The module contains both *YAO* and *NUC1*, and *NUC1* is the most highly expressed gene within the module. **b**, Structure of the pHY07 vector for CRISPR/Cas9 gene editing and the sgRNA target site within *GL2*. In pHY07, the *NUC1* promoter is used to drive *Cas9* expression. **c**, T0 seeds observed under a fluorescent microscope. Positive transgenic seeds emitting fluorescence are visually distinct from other seeds. **d**, Phenotypes of T1 generation plants transformed with the pHY07-GL2 construct. Three categories of plants are shown: WT-like, chimera, and glabrous. Scale bar corresponds to 1 cm. **e**, Mutation efficiency in T1 transgenic plants transformed with pHY07-GL2 or CRISPR/Cas9 vectors driven by other promoters, as determined by Sanger sequencing (n = 39/42/46/35/39). Homo, homozygous; Bi/He, bi-allelic or heterozygous; Chi: chimeric; WT: wild type.

### Gene editing efficiency in T1 generation

In order to evaluate its gene editing efficiency, we used pHY07 to target the *GLABRA2* (*GL2, AT1G79840*) gene of Arabidopsis. This gene regulates leaf trichome development, and its mutation results in a glabrous phenotype due to defects in leaf trichomes [24]. Because this phenotype is easy to observe, it was often selected as a target gene for testing gene editing efficiency [13, 25]. In our study, we used a single sgRNA driven by the *AtU6* promoter to target *GL2*, which was cloned into pHY07 to generate the final CRIPSR vector pHY07-GL2 (Fig. 1b). The vector was used to transform Arabidopsis Col-0 plants via the *Agrobacterium*-mediated floral dip method [11]. The positive transformed T0 seeds constitutively expressed mCherry and emitted red fluorescence under a fluorescence microscope, which was visually distinct from other negative seeds (Fig. 1c). The fluorescent seeds were selected and subjected to a second screening on 1/2 MS agar plates containing 15 μg/mL hygromycin, and 42 positive T1 plants were obtained and planted in the soil. After two weeks, based on their leaf surface phenotypes, we divided these plants into three categories: 19 (45%) with a glabrous phenotype where all leave surfaces are uniformly hairless, 15 (36%) with a chimera phenotype where portions of the leave surfaces containing no hair, and 8 (19%) with a wild-type phenotype (Fig. 1d and Supplementary Table 1). In order to more accurately detect and quantify gene editing efficiency, we isolated genomic DNA from these plants’ leaves, amplified the target site by PCR, and subjected them to Sanger sequencing. The resulted sequence chromatograms showed that 15 (36%) of the T1 plants appeared to contain homozygous mutations, 8 (19%) contained bi-allelic/heterozygous (bi-allelic or heterozygous) mutations, 18 (43%) contained chimeric mutations, and 1 (2.4%) was WT (Fig. 1e). The overall mutation efficiency was 97.6%, while 55% of the T1 plants harbored apparently heritable nonchimeric (homozygous/bi-allelic/heterozygous) mutations. Interestingly, we found that 3 plants with WT-like phenotypes were indeed homozygous mutants. Among them, two have 9 bp deletions while the other has 27 bp deletion near the sgRNA target site. These mutations just affected a limited number of amino acids, which might not interfere the function of GL2. To examine the specificity of the gene editing, we identified three potential off-target sites for the *GL2* sgRNA. We sequenced these off-target sites from 10 independent T1 plants, and found no mutations, thus proving the high specificity of our pHY07 CRISPR/Cas9 system.

Besides *NUC1*, there are already a handful of alternative promoters that have been used to drive *Cas9* expression for gene editing efficiency improvement. We selected three representative promoters, together with the constitutive CaMV 35S promoter (*p35S*), to compare their efficiency with the *NUC1* promoter. We replaced the *NUC1* promoter within pHY07 with the *p35S, EC1*.*2en-EC1*.*1 fusion, YAO*, or *CDC45* promoters, and used the modified systems to target *GL2* via the same sgRNA. None of the T1 plants transformed with p35S-Cas9-GL2 showed any phenotype. Among the T1 plants transformed with pEC1.2en-EC1.1-Cas9-GL2, pYao-Cas9-GL2 and pCDC45-Cas9-GL2, 13%, 8.6%, and 10% showed a uniform glabrous phenotype, and 6.5%, 83%, and 90% showed a chimera phenotype, respectively (Supplementary Table S1). Sequencing of the sgRNA target site revealed that none of T1 plants transformed with p35S-Cas9-GL2 contained mutations, while 23%-28% of the T1 plants transformed with pEC1.2en-EC1.1-Cas9-GL2, pYao-Cas9-GL2, and pCDC45-Cas9-GL2 contained apparently nonchimeric mutations (Fig. 1e). The results indicated that the nonchimeric mutation efficiency of the CRISPR/Cas9 system driven by the *NUC1* promoter is much higher than that of other promoters.

### Heritability of the mutations

An important question regarding CRISPR/Cas9 gene editing system is whether the mutations observed in T1 generation can be stably inherited into the next generation. We further characterized the mutation statuses of the T2 plants from the T1 plants transformed with the pHY07-GL2 vector. Since for gene editing experiments, it is preferred to have Cas9-free stable mutants, we focused on Cas9-free T2 plants only. We randomly chose 10 independent T1 lines for the characterization. From each of these lines, following a procedure reported before [26], we selected 10 non-fluorescent and Cas9-free seeds for planting (Fig. 2a). Because some homozygous mutants displayed WT-like phenotypes in the T1 generation, the *GL2* loss of function phenotype might not be an accurate indicator of overall mutation efficiency. We directly used Sanger sequencing to genotype the PCR-amplified *GL2* target site from the T2 plants. Among the 100 Cas9-free T2 plants tested, 66 harbored homozygous mutations, 33 contained bi-allelic/heterozygous mutations, and 1 was WT (Fig. 2b). Thus, 99% of them harbored mutations that were stably inherited from the T1 generation, since they did not contain the Cas9 cassette to generate new mutations. Interestingly, 6 of the 10 selected T1 plants were chimeric mutants, but 98% of their Cas9-free T2 offspring had inherited mutations, indicating that chimeric mutants generated by pHY07 also had very high chances to pass on heritable mutations to the next generation. On the other hand, we observed 3 bi-allelic/heterozygous mutants among the 20 Cas9-free T2 offspring from 2 homozygous T1 mutants. These T1 homozygous mutants might contained additional minor mutant alleles that cannot be detected from Sanger sequencing chromatograms. Thus, when targeting the *GL2* gene with pHY07, the Cas9-free T2 plants had a very high chance to inherit stable mutations from their T1 parents, which greatly speeded up the process to obtain Cas9-free stable mutants.

**Figure 2.**
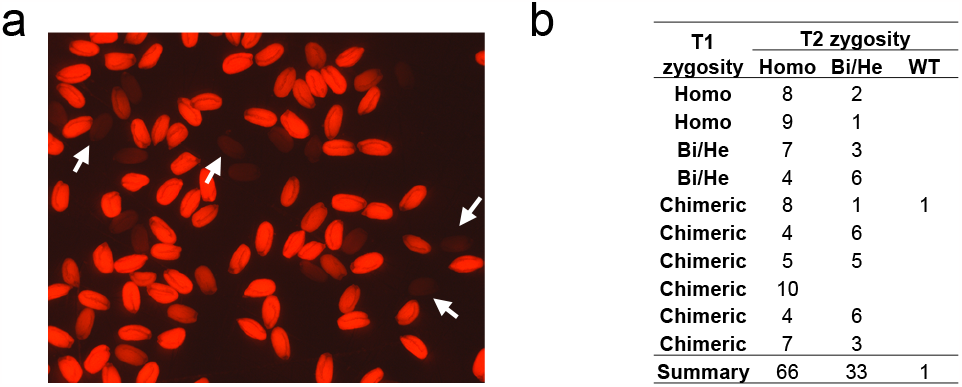
Mutation rate in T2 generation Cas9-free Arabidopsis plants. **a**, Seeds from T1 generation transgenic plants transformed with pHY07-GL2 observed under a fluorescent microscope. Non-fluorescent seeds (white arrows) were selected and planted to obtain Cas9-free T2 plants. **b**, Mutation status of the Cas9-free T2 plants. Ten independent T1 lines were randomly chosen, and from each of these lines 10 non-fluorescent seeds were selected for planting. The zygosity of the T2 plants were determined by Sanger sequencing. Homo, homozygous; Bi/He, bi-allelic or heterozygous; Chimeric: chimeric; WT: wild type.

### Gene editing efficiency on other genes

To further evaluate its gene editing efficiency, we also used pHY07 to target a pair of homologous transcription factor genes *TRY* and *CPC* via a single sgRNA that has been validated before [14]. *TRY* and *CPC* are negative regulators of trichome development, and their knockout plants displayed an increased number of trichomes in the leaves [14]. Forty positive seedlings were selected for the T1 generation, and 100% of them showed a completely mutant phenotype (Fig. 3a). In order to evaluate the heritability of the mutations, we randomly chose 10 independent T1 plants, and selected 10 non-fluorescent Cas9-free seeds from each of these lines for planting. We then PCR amplified and genotyped the target sites within the *TRY* and *CPC* genes from these Cas9-free T2 plants. The results showed that for the *TRY* and *CPC* genes, 68% and 59% of the plants contained homozygous mutations, and 26% and 35% contained bi-allelic/heterozygous mutations respectively (Fig. 3b). The overall mutation rates in Cas9-free T2 plants are 94% for both genes.

**Figure 3.**
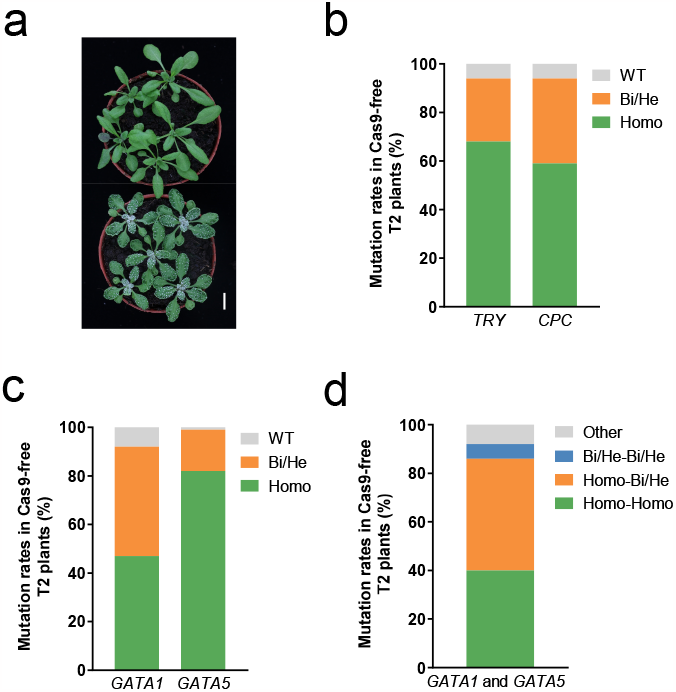
Editing efficiency of pHY07 on additional genes. **a, b**, Gene editing efficiency on *TRY* and *CPC*. A single sgRNA was used to edit both *TRY* and *CPC*. Forty T1 transgenic plants were grown and all displayed *TRY*/*CPC* knock-out phenotypes. Shown in **(a)** are 5 WT plants (top) and 5 T1 transgenic plants (bottom). Scale bar corresponds to 1 cm. Shown in **(b)** are *TRY* and *CPC* mutation status in the Cas9-free T2 plants (n = 100). **c, d**, Multiplex gene editing efficiency on *GATA1* and *GATA5* using pHY07. Mutation status of individual *GATA1* and *GATA5* genes **(c)** and their joint mutations status **(d)** in Cas9-free T2 plants (n = 100) are shown. In both **(a, b)** and **(c, d)**, 10 independent T1 lines were randomly chosen, and from each of these lines 10 non-fluorescent seeds were selected for planting to obtain Cas9-free T2 plants. The zygosity of the T2 plants were determined by Sanger sequencing. Homo, homozygous; Bi/He, bi-allelic or heterozygous; WT: wild type; Homo-Homo, both genes with homozygous mutations; Homo-Bi/He, one gene with homozygous mutation and the other with bi-allelic/heterozygous mutation; Bi/He-Bi/He, both genes with bi-allelic/heterozygous mutations; Other, other mutation types.

### Multiplex gene editing efficiency

We also tested the efficiency of pHY07 in multiplex gene editing. We chose two uncharacterized genes *GATA1* and *GATA5* from the GATA transcription factor family. Two sgRNAs, targeting *GATA1* and *GATA5* and driven by the *AtU6* and *AtU3* promoters respectively, were cloned into a single pHY07 vector and used to transform Arabidopsis Col-0 plants. Thirty-eight positive transformed T1 plants were planted, but they did not show any obvious phenotypes under normal growth conditions. We then randomly picked 10 independent T1 plants and selected from each 10 non-fluorescent Cas9-free seeds for planting. Among the 100 T2 plants tested, for *GATA1*, 47 and 45 harbored homozygous and bi-allelic/heterozygous mutations respectively; and for *GATA5*, 82 and 17 contained the same two types of mutations (Fig. 3c). Importantly, we identified 40 T2 plants with homozygous mutations on both genes, 46 with homozygous mutation on one gene and bi-allelic/heterozygous mutation on the other, and 6 with bi-allelic/heterozygous mutations on both genes (Fig. 3d). Thus, we obtained stable inherited mutations on both genes within 92% of the Cas9-free T2 plants, indicating that our pHY07 system also has very high efficiency when editing multiple target sites simultaneously.

### Other genes from the same modules as alternative promoters

Beside *NUC1*, we also tested other highly expressed promoters within Module #21 to evaluate their potential usage on gene editing. Three genes, *At2G39390, At3G03920*, and *At5G67510*, were chosen and their promoters were used to drive *Cas9* expression to construct three modified CRISPR/Cas9 systems, pHY14, pHY15, and pHY16. We used these systems to edit *GL2* via the same sgRNA as used before. Among their positive T1 transformed plants, 16%, 22%, and 52% showed a uniform glabrous phenotype (Supplementary Table 1). Genotyping of the PCR amplified target sites indicated 32%, 11%, 45% of them have apparently nonchimeric mutations (Fig. 4). The results indicated that these promoters also function similarly to *NUC1*, although their efficiencies are a little bit lower. The applicability of *NUC1* and these three alternative promoters validated our data-driven approach to identify promoters for improving the efficiency of CRISPR/Cas9-based gene editing.

**Figure 4.**
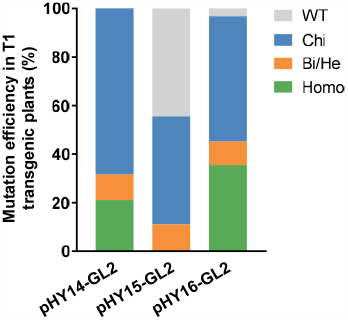
Three alternative promoters for driving *Cas9* expression. Three other highly expressed genes were identified from Module #21 of AtGGM2014. Their promoter sequences were used as alternative promoters to drive *Cas9* expression in three modified CRISPR/Cas9 systems: pHY14: *pAt2G39390-Cas9*; pHY15: *pAt3G03920-Cas9*, and pHY16: *pAt5G67510-Cas9*. Shown here is the gene editing efficiency on *GL2* in T1 generation transgenic plants (n = 19/9/31) using these modified systems. Homo, homozygous; Bi/He, bi-allelic or heterozygous; Chi: chimeric; WT: wild type.

## Discussion

In the current study, we employed a data-driven approach to improve genome editing efficiency of the CRISPR/Cas9 system in Arabidopsis. Previous works have shown that the promoter driving *Cas9* expression plays critical role in determining the efficiency of heritable mutations. Our gene co-expression network analysis identified the *NUC1* promoter as an ideal candidate to drive the expression of *Cas9* for optimizing the editing system. *NUC1* was contained within the same gene co-expression module as *YAO*, indicating it should drive the expression of *Cas9* in similar tissues as *YAO* and opt for generating heritable mutations. It should also deliver the highest editing efficiency since it was the most highly expressed gene within the module. We then generated a modified CRISPR/Cas9 system pHY07 using the *NUC1* promoter to drive Cas9 expression. Using the *GL2* gene as an example, our results showed that the editing efficiency of Cas9 driven by the *NUC1* promoter was indeed much higher than other promoters used before (Fig. 1e). Of the 42 positive transformed T1 plants transformed with pHY07-GL2, 55% contained apparently nonchimeric mutations, while for other promoters the nonchimeric mutation rates are between 0 – 28%. We also tested another three potential promoters from the same gene co-expression module, which resulted in nonchimeric mutation rates of 22 – 45% in T1 generation, further validating our data-driven approach for enhancing CIRSPR/Cas9 gene editing efficiency.

Our optimized CRISPR/Cas9 system pHY07 makes it easy to obtain Cas9-free stably transmissible mutations in Arabidopsis. Gene editing in Arabidopsis requires the insertion of the CRISPR/Cas9 cassette into the plant’s genome. After the target site(s) has been edited, it is often advisable to segregate out this cassette before downstream functional characterization. We incorporated a *mCherry* gene driven by a seed specific promoter *At2S3* in pHY07 for visually selecting Cas9-contained T0 seeds and Cas9-free T1 seeds [19, 20]. Coupling the high gene editing efficiency brought by the *NUC1* promoter and the fluorescence selection via mCheery, pHY07 speeds up the process to obtain Cas9-free stable mutants in T2 Arabidopsis plants. For example, when targeting the *GL2* or the *TRY* and *CPC* genes, we found stable mutations inherited from the previous generation in 94 – 99% of the Cas9-free T2 plants. Similar efficiency was also observed when editing two genes *GATA1* and *GATA5* simultaneously, indicating our system also works for multiplex editing. Thus, we believe our system will become a valuable tool for Arabidopsis gene function studies.

In conclusion, we have optimized and constructed a highly efficient and convenient CRISPR/Cas9 system for gene editing in Arabidopsis. Our data-driven approach based on gene co-expression network analysis can also be used to select the appropriate promoter sequences in other scenario, e.g., choosing promoters to over-express a gene while keeping its expression patterns or to over-express a gene in a tissue or condition-specific manner.

## Methods

### Candidate gene promoter identification

We used a published Arabidopsis gene co-expression network AtGGM2014 to identify co-expression modules containing the genes whose promoters have been used before for driving *Cas9* expression [21]. The *Yao* gene was found to be within Module #21, which contains 108 genes in total. To compare the expression levels of these genes, we downloaded the raw data of all publically available Arabidopsis RNA-Seq transcriptomes from NCBI’s Sequence Read Archive database as of December 2018. After removing RNA-Seq runs for small RNAs, the remaining runs were trimmed with Trimmomatic (version 0.36) and mapped onto the Arabidopsis genome (Araport11) using STAR 34 (version 020201) [27, 28]. The resulted bam files were used to calculate the transcripts per million (TPM) gene expression values via RSEM (v1.3.0) [29]. We further filtered out the runs meeting any of these criteria: total reads number < 1,000,000; unique mapping rate < 50%; total mapping rate < 70%; or having less than 5,000 genes with TPM ≥ 1. The expression profiles from the remaining runs were then log-transformed via log_2_(TPM+1) and used to calculate the average expression levels for the genes within Module #21. The gene with highest average expression level was chosen as a candidate gene promoter for improving CRISPR/Cas9 gene editing efficiency.

### Cas9 plasmids construction

All plasmids were constructed via recombination using ClonExpress ® II One Step Cloning Kit (Vazyme, Nanjing, China). The *zCas9* sequence, together with the *Pisum sativum rbcS* E9 terminator sequence (rbcS_E9t), was amplified from the pHEE401 vector [14], and recombined into the pCambia1300 binary vector digested with *EcoR*I to obtain pCambia1300-Cas9. A DNA fragment containing the *mCherry* fluorescent protein gene driven by the *At2S3* promoter [19, 20] was synthesized and recombined into pCambia1300-Cas9 through *EcoR*I and *Hind*III restriction sites to obtain pCambia1300-Cas9-mCherry. A 1,472 bp *NUC1* gene promoter fragment was amplified from the Arabidopsis Col-0 genomic DNA and recombined into pCambia1300-Cas9-mCherry through the *Nco*I restriction site, to get pCambia1300-pNUC1-Cas9-mCherry, which was named as pHY07. The CRISPR/Cas9 plasmids with *Cas9* driven by other promoters were also constructed via cloning and recombining the corresponding promoter sequence into pCambia1300-Cas9-mCherry.

### Guide RNA design and sgRNA plasmids construction

To facilitate the assembly of sgRNA cassettes, we synthesized a DNA fragment containing AtU6-26p promoter, gRNA scaffold, and AtU6-26t terminator, and ligated it into the pUC57 plasmid through *Afl*III and *Aat*II restriction sites to obtain the pUC57_U6 plasmid. Another plasmid pUC57_U3, containing the gRNA scaffold, the 67 bp AtU6 terminator and AtU3b promoter sequence, was also constructed. To obtain sgRNA cassette with one sgRNA, pUC57_U6 was used as template to conduct two PCR amplifications with one of the primers containing the sgRNA sequence, and the 2 resulted PCR fragments were recombined to obtain pUC57_1sgRNA intermediate vector. To obtain sgRNA cassette with two sgRNAs, the pUC57_U6 and pUC57_U3 plasmids were used as templates to conduct two PCR amplifications, with primers containing the two sgRNAs’ sequences and their reverse complementary sequences. The 2 resulted PCR fragments were recombined to obtain pUC57_2sgRNA intermediate vector. The pUC57_1sgRNA vector contains a sgRNA cassette to expression 1 sgRNA driven by *AtU6* promoter, and the pUC57_2sgRNA vector contains a sgRNA cassette to express 2 sgRNAs driven by *AtU6* and *AtU3* promoters respectively. The sgRNA cassette was then amplified via PCR and recombined into pHY07 via the *Hind*III restriction site to obtain the final binary vector for plant transformation. A detailed procedure can be found in Supplemental Method 1.

We designed sgRNAs targeting *GL2, GATA1*, and *GATA5* via CRISPR-P 2.0, and obtain a sgRNA targeting both *TRY* and *CPC* from a previous report [14]. The sgRNAs used were: *GL2*: GGCTTGATGCAGCAATGGCG; *GATA1*: ACGAACATACGCAACCACCG; *GATA5*: GCCTGACGCTGCTCTTCAACG; *TRY*/*CPC*: GAATATCTCTCTATCTCCTC. The first base of these sgRNAs are not in the target site but serves as the transcription start site for *AtU6* or *AtU3* promoters. The sgRNAs targeting *GL2*, or *TRY* and *CPC* were cloned into pHY07 to generate pHY07-GL2 or pHY07-TRY/CPC. The sgRNAs targeting *GATA1* and *GATA5* were cloned into pHY07 simultaneously to generate pHY07-GATA1-GATA5. The primers used are listed in Supplementary Table 2.

### Arabidopsis growth conditions

Surface-sterilized Arabidopsis seeds were stratified at 4 °C in dark for 2 days and placed on vertical 1/2 MS (containing 15μg/mL hygromycin) plates supplemented with 1% (w/v) sucrose and 0.8% (w/v) agar at 22 °C under 16 h day/ 8 h night cycle for growing. After 7 days, seedlings with long roots were potted in soil for further growing in a growth chamber at 22 °C under a 16 h day/8 h night photoperiod.

### Arabidopsis transformation and selection of positive plants

Arabidopsis Columbia (Col-0) plants were transformed with Agrobacterium strain GV3101 by the floral dip method [11]. After harvesting the T0 generation seeds, positive transgenic seeds emitting mCherry florescence were selected under a stereoscopic fluorescence microscope (Axio Zoom.V16, Zeiss). The selected seeds were subjected to a second screening by germinating on 1/2MS agar medium containing 15μg/mL hygromycin before being transferred to soil for growing. The seeds harvested from the T1 generation plants were also used to selected non-fluorescent Cas9-free seeds for further characterizations.

### Mutation detection

The CTAB method extracted from rosette leaves of transgenic Arabidopsis genomic DNA[30]. PCR amplified genomic regions flanking the target sites using primer pairs listed in Supplementary Table 2. Mutations in target genes were detected by sequencing of the PCR products and the pattern of zygosity was identified by analyzing the resulted Sanger sequence chromatograms.

## Acknowledgements

This work was supported by grants from the National Natural Science Foundation of China (31770268), the Strategic Priority Research Program of the Chinese Academy of Sciences (XDA24010303), the Fundamental Research Funds for the Central Universities (WK2070000091), and University of Science and Technology of China (Start-up fund to S.M.).

## Author contributions

SM designed and supervised the project. HG, YW, YX, YZ, EH, YP, ZG, YL, and YQ performed the experiments and analyses. HG, YW, and SM wrote the manuscript. All authors reviewed and approved the final manuscript.

## Supplementary Information

**Supplementary Table 1**. Phenotypic characterization of T1 generation Arabidopsis.

**Supplementary Table 2**. List of primers used in the study.

**Supplementary Method 1**. The procedure to clone 1 or 2 sgRNA(s) into pHY07.

