## Supplementary Information for "Data-driven optimization yielded a highly-efficient CRISPR/Cas9 system for gene editing in Arabidopsis"

The following Supporting Information is available for this article:

**Supplementary Table 1.** Phenotypic characterization of T1 generation Arabidopsis.

**Supplementary Table 2.** List of primers used in this study.

**Supplementary Method 1.** The procedure to clone 1 or 2 sgRNA(s) into pHY07.

**Supplementary Table 1.** Phenotypic characterization of T1 generation Arabidopsis

| | **Phenotype of T1 generation transgenic plants** | | | |
|---|---|---|---|---|
| | **NO. of uniform glabrous plants** | **NO. of chimera plants** | **NO. of wild-type like plants** | **Total** |
| **p35S-Cas9-GL2** | 0 | 0 | 39 (100%) | 39 |
| **pHY07(pNUC1-Cas9)-GL2** | 19 (45%) | 15 (36%) | 8 (19%) | 42 |
| **pEC1.2en-EC1.1-Cas9-GL2** | 6 (13%) | 3 (6.5%) | 37 (80%) | 46 |
| **pYAO-Cas9-GL2** | 3 (8.6%) | 29 (83%) | 3 (9%) | 35 |
| **pCDC45-Cas9-GL2** | 4 (10%) | 35 (90%) | 0 | 39 |
| **pHY14(pAT2G39390-Cas9)-GL2** | 3 (16%) | 13 (68%) | 3 (16%) | 19 |
| **pHY15(pAT3G03920-Cas9)-GL2** | 2 (22%) | 2 (22%) | 5 (56%) | 9 |
| **pHY16(pAT5G67510-Cas9)-GL2** | 16 (52%) | 13 (42%) | 2 (6%) | 31 |

**Supplementary Table 2.** List of primers used in this study

| | |
|---|---|
| **Primers for vector construction** | |
| Cas9-F | GTACCGAGCTCGAATTCCATGGATTACAAGGACCACGACGGGGATTACAAGGACCACGACATTGATT |
| Cas9-R | TATGACCATGATTACGAATTGTTGTCAATCAATTGGCAAG |
| mCherry-F | CGACGGCCAGTGCCAAGCTAAGCTGGCACAACTATATTTC |
| mCherry-R | CGACCTGCAGGCATGCAAGCTTGATAATTTATTTGAAAATTC |
| pNUC1-F | ACCGAGCTCGAATTCCATGACTGCTCGCTGCTTTGTAGAATG |
| pNUC1-R | CGTGGTCCTTGTAATCCATGGAGAACTGAGAAAGAGACGACTG |
| p35S-F | ACCGAGCTCGAATTCCATGTGAGACTTTTCAACAAAGGGTA |
| p35S-R | CGTGGTCCTTGTAATCCATGGTGTTCTCTCCAAATGAAATGAAC |
| pEC1.1-1.2-F | ACCGAGCTCGAATTCCATGGTGAATAAAAGCATTTGCGTTTG |
| EC1.2R | AGCTAATTCATGATAGGCGTTACTAGCTTAGTGGTGATTTAAGAGT |
| EC1.1F | ACTCTTAAATCACCACTAAGCTAGTAACGCCTATCATGAATTAGCT |
| pEC1.1-1.2-R | CGTGGTCCTTGTAATCCATGGTTCTCAACAGATTGATAAGGTC |
| pYAO-F | ACCGAGCTCGAATTCCATGGATGGGAAATTCATTGAAAACC |
| pYAO-R | CGTGGTCCTTGTAATCCATGGCTCCTTTCTTCTTCTCGTTGT |
| pCDC45-F | ACCGAGCTCGAATTCCATGCTCCTGATGATAAAGGTGGGA |
| pCDC45-R | CGTGGTCCTTGTAATCCATGGTTCCGTGAAATTGAATCACCC |
| pAT2G39390-F | CCGGGTACCGAGCTCGAATTCGATCCTCTGTTTTTTTGTCCTG |
| pAT2G39390-R | CGTGGTCCTTGTAATCCATGGTGCGTCTTCGAGCTCTCTCTA |
| pAT3G03920-F | CCGGGTACCGAGCTCGAATTCTAATGAAACATGGAGCATATTCT |
| pAT3G03920-R | CGTGGTCCTTGTAATCCATGGCTTTTCTCCTTTCACAGATTCC |
| pAT5G67510-F | CCGGGTACCGAGCTCGAATTCCATCCATATAAAAAAAATTCGAAAG |

| | |
|---|---|
| pAT5G67510-R | CGTGGTCCTTGTAATCCATGGCGCCGCTGAATTTGTGAAGAA |
| **Primers for sgRNAs cloning** | |
| p57-U6-GL2-P1F | GTCGAAGTAGTGATTGGCTTGATGCAGCAATGGCGGTTTTAGAGCTAGAAATAGC |
| p57-U6-P2R | GAGTAAACTTGGTCTGACAG |
| p57-U6-P3F | CTGTCAGACCAAGTTTACTC |
| p57-U6-P4R | AATCACTACTTCGACTCTAGC |
| p57-U6-TRY/CPC-P1F | GTCGAAGTAGTGATTGAATATCTCTCTATCTCCTCGTTTTAGAGCTAGAAATAGC |
| GATA5-PCR1-F | CCTGACGCTGCTCTTCAACGGTTTTAGAGCTAGAAATAGCAAG |
| GATA1-PCR1-R | CGGTGGTTGCGTATGTTCGTGACCAATGGTGCTCCCTCAGTG |
| GATA1-PCR2-F | CGAACATACGCAACCACCGGTTTTAGAGCTAGAAATAGCAAG |
| GATA5-PCR2-R | CGTTGAAGAGCAGCGTCAGGCAATCACTACTTCGACTCTAGC |
| pUC57-seq-F | CCAGGATTAGAATGATTAGGC |
| pHY07-Fu-F | TTTCAAATAAATTATCAAGCTACCACGTAGGATCCGCTAGC |
| pHY07-Fu-R | CGACCTGCAGGCATGCAAGCTCTTTTGCTGGCCTTTTGCTC |
| MAS-seq-F | TCAGTAATCTCGGCCAATATC |
| **Primers for amplifying and genotyping the sgRNA target sites** | |
| GL2-F | GCTTCTCTTCTGAAATGTCGA |
| GL2-R | AAGAGCTGGAGAAACAGGTAA |
| GATA1-F | CGTTACACTTGATTGTGGCTT |
| GATA1-R | TCAAAGGGTAGCTTCGTCTTT |
| GATA5-F | TGTTTGTGGGTCGTTTTCGTA |
| GATA5-R | GGAAGAGAGAGTTCGCTTGTA |
| TRY-F | ACTATTCTTATCAGTACGTACTC |
| TRY-R | CAAGAACCACTACTATACATCTA |
| CPC-F | TGTCAGAACTCACTTTGGCTA |
| CPC-R | CACAGCTACTAATACTATACTTC |

**Supplementary Method 1.** The procedure to clone 1 or 2 sgRNA(s) into pHY07

**Materials:**

Backbone vectors: pUC57_U6 (Amp$^+$); pUC57_U3 (Amp$^+$); pHY07 (Kan$^+$).

Restriction enzyme: *Hind*III

ClonExpress II One Step Cloning Kit (Catalog #C112, Vazyme, Nanjing, China)

sgRNA design: CRISPR-P (http://crispr.hzau.edu.cn/cgi-bin/CRISPR2/CRISPR)

**Methods:**

### A. Clone a single sgRNA into pUC57_U6 through recombination

1. Design a sgRNA to target a gene of interest.
   The sgRNA will be driven by U6 promoter, and its transcription starts with G. Therefore, if the first base of the sgRNA is "G", the G can be omitted since it's already in the primer.

2. Perform two PCRs, using pUC57_U6 as template:
   1) Conduct the first PCR to generate Fragment 1

      Forward primer: **p57-U6-P1F**
      GTCGAAGTAGTGATTGNNNNNNNNNNNNNNNNNNNGTTTTAGAGCTAGAAATAGC

      Replace NNNNNNNNNNNNNNNNNNN with the actual sgRNA sequence.

      Reverse primer: **p57-U6-P2R**

      GAGTAAACTTGGTCTGACAG

      Expected fragment size: 1351bp.

   2) Conduct the second PCR to generate Fragment 2

      Forward primer: **p57-U6-P3F** CTGTCAGACCAAGTTTACTC

      Reverse primer: **p57-U6-P4R** AATCACTACTTCGACTCTAGC

      Expected fragment size: 1270 bp.

3. Gel purify the two fragments using Gel Extraction Kit and elute in EB solution to get rid of the pUC57_U6 vector (Important!!! pUC57_U6 has the same antibiotic resistance as the end vector).

4. Use ClonExpress II One Step Cloning Kit to recombine the two fragments and generate a vector with 1 sgRNA, named as pUC57_1sgRNA.
   Set up recombination reaction and incubate at 37°C for 30 min:

| | |
|---|---|
| Fragment 1 | X µl |
| Fragment 2 | Y µl |
| 5 × CE II Buffer | 4 µl |
| Exnase II | 2 µl |
| Add dd$H_2O$ up to 20 µl | |

5. Transform into *E. coli* DH5α competent cells and grow on Agar plates supplemented with ampicillin. Purify plasmids from the positive clones, and verify the plasmids by sequencing:

   Sequencing primer: **pUC57-seq-F** CCAGGATTAGAATGATTAGGC

### B. Clone two sgRNAs into pUC57_U6 through recombination

1. Design two sgRNAs to target two different genes.
   sgRNA1 will be driven by U6 promoter, and its transcription starts with G. sgRNA2 will be driven by U3 promoter, and its transcription starts with A. Therefore, if the first base of sgRNA 1 is "G",

the G can be omitted since it's already in the primer. And if the first base on sgRNA 2 is "A", the A can be omitted since it's already in the primer.

2. Conduct first PCR to generate Fragment 1. Purify the fragment ( ~480 – 490 bp) after PCR to get rid of the pUC57_U3 vector (Important!!! pUC57_U3 has the same antibiotic resistance as the end vector).
   Fragment 1:

   Use pUC57_U3 as back-bone template to do the PCR reaction.

   Forward primer: **sgRNA1-PCR1-F**
   NNNNNNNNNNNNNNNNNNNNGTTTTAGAGCTAGAAATAGCAAG

   Replace NNNNNNNNNNNNNNNNNNNN with the sequence of sgRNA1.

   Reverse primer: **sgRNA2-PCR1-R**
   NNNNNNNNNNNNNNNNNNNNTGACCAATGGTGCTCCCTCAGTG

   Replace NNNNNNNNNNNNNNNNNNNN with the reverse complement sequence of sgRNA2.

3. Conduct second PCR to generate Fragment 2. Purify the fragment (~2600 bp) after PCR to get rid of the pUC57_U6 vector (Important!!! pUC57_U6 has the same antibiotic resistance as the end vector). (Note: pUC57_U6 can also be digested with *Pme*I and linearized first, and then used for PCR.)
   Fragment 2:

   Use pUC57_U6 as back-bone template to do the PCR reaction.

   Forward primer: **sgRNA2-PCR2-F**
   NNNNNNNNNNNNNNNNNNNNGTTTTAGAGCTAGAAATAGCAAG

   Replace NNNNNNNNNNNNNNNNNNNN with the sequence of sgRNA2.

   Reverse primer: **sgRNA1-PCR2-R**
   NNNNNNNNNNNNNNNNNNNNCAATCACTACTTCGACTCTAGC

   Replace NNNNNNNNNNNNNNNNNNNN with the reverse complement sequence of sgRNA1.

4. Use ClonExpress II One Step Cloning Kit to recombine the two fragments and generate a vector with 2 sgRNAs, named as pUC57_2sgRNA.
   Set up recombination reaction and incubate at 37°C for 30 min:

| | |
|---|---|
| Fragment 1 (step 2) | X µl |
| Fragment 2 (step 3) | Y µl |
| 5 × CE II Buffer | 4 µl |
| Exnase II | 2 µl |
| Add dd$H_2O$ up to 20 µl | |

5. Transform into *E. coli* DH5α competent cells and grow on Agar plates supplemented with ampicillin. Purify plasmids from the positive clones, and verify the plasmids by sequencing. Then clone the sgRNA cassette into the final pHY07 vector.

   Sequencing primer: **pUC57-seq-F** CCAGGATTAGAATGATTAGGC

### C. Clone the sgRNA cassette into the binary vector pHY07

1. Amplify the sgRNA-cassette by PCR from pUC57_1sgRNA or pUC57_2sgRNA, and purify the PCR fragment. Expected fragment size: 1sgRNA~826bp, 2sgRNA~1287bp.

   Use pUC57_1sgRNA or pUC57_2sgRNA as template.
   Forward primer: **pHY07-Fu-F**

   TTTCAAATAAATTATCAAGCTACCACGTAGGATCCGCTAGC

   Reverse primer: **pHY07-Fu-R**

   CGACCTGCAGGCATGCAAGCTCTTTTGCTGGCCTTTTGCTC

2. Digest 1μg of pHY07 with *Hind*III and purified the linearized vector fragment.

3. Set up recombination reaction and incubate at 37°C for 30 min:

| | |
|---|---|
| PCR fragment (step 1) | X μl |
| Vector fragment (step 2) | Y μl |
| 5 × CE II Buffer | 4 μl |
| Exnase II | 2 μl |
| Add dd$H_2O$ up to 20 μl | |

4. Transform into *E. coli* DH5α competent cells and grow on Agar plates supplemented with kanamycin. Purify plasmids from the positive clones, and verify the plasmids by sequencing.

   Sequencing primer: **MAS-seq-F** TCAGTAATCTCGGCCAATATC
